# Wave-inspired MEW scaffolds for enhanced ligament tissue regeneration

**DOI:** 10.64898/2025.12.21.695497

**Authors:** Joanna Babilotte, Clarissa Tomasina, Alberto Sensini, Tim ten Brink, Pieter Emans, Monize C. Decarli, Paul Wieringa, Lorenzo Moroni

## Abstract

Ligament injuries remain a major clinical challenge due to the limited intrinsic healing capacity of these fibrous tissues. Here, we demonstrate the use of melt electrowriting (MEW) to fabricate poly(ε-caprolactone) (PCL) scaffolds with precisely engineered wave architectures that mimic the hierarchical organization and nonlinear mechanics of native ligaments. By tuning fiber geometry, we achieved scaffolds with distinct mechanical behaviors ranging from highly compliant to structurally resilient, enabling architecture-driven modulation of elastic modulus and fatigue response. Mechanical testing revealed that wave-patterned scaffolds dissipate energy efficiently and adapt structurally under cyclic loading, reproducing key features of ligament-like viscoelasticity. When cultured with human anterior cruciate ligament (ACL) cells, the scaffolds supported adhesion, proliferation, and spatially organized alignment, together with the expression of ligament-associated markers. The results demonstrate that MEW scaffolds provide a favorable environment for ligament cell adhesion and matrix synthesis, while highlighting the strong influence of geometry on cell organization and early matrix production. Overall, this study establishes wave-based MEW architectures as a versatile platform to guide ligament tissue formation.

## Introduction

Ligaments are essential components of the musculoskeletal system, connecting bones across joints to provide mechanical stability and guide motion during daily activities. Because they operate under continuous load, they are frequently subjected to excessive strain or rupture, leading to joint instability and long recovery periods. Ligament injuries are among the most common musculoskeletal disorders and represent a major socio-economic burden due to the need for surgical repair and extensive rehabilitation [1]. In the Netherlands, early anterior cruciate ligament (ACL) reconstruction in adults aged 18–65 incurs societal costs of €48,460– €78,179 per quality-adjusted life year gained, underscoring the high economic impact of treatment decisions [2].

Ligaments have a limited intrinsic healing capacity owing to their low vascularization and cellularity [3]. As a result, spontaneous repair is often incomplete, leading to scar tissue formation and inferior mechanical properties [4,5]. Surgical reconstruction using autografts or allografts remains the current gold standard but is associated with several drawbacks, including donor-site morbidity, poor graft integration, and a high risk of re-injury or long-term joint degeneration [4,6,7]. Synthetic grafts have also been explored, yet they often fail to provide sufficient biocompatibility and long-term mechanical stability [4,6]. Integration of grafts with the host tissue remains a major clinical challenge, often leading to incomplete healing and reinjury [3]. These challenges highlight the pressing need for new regenerative approaches capable of restoring both the structure and function of native ligament tissue.

Ligaments are dense connective tissues composed mainly of type I collagen arranged in a highly organized hierarchical architecture. The collagen fibrils are bundled into fibers displaying a characteristic crimped or wavy pattern [8–10]. This crimp enables the ligament to stretch and uncrimp under small loads, a behavior reflected in the initial non-linear portion of the stress– strain curve, known as the “toe region” [10]. Once the crimp is straightened, the fibers bear the load and provide high tensile strength [10]. This hierarchical structure imparts unique mechanical behavior, combining flexibility, resilience, and strength [10,11]. It also provides structural cues that guide fibroblast alignment and extracellular matrix (ECM) deposition along the load-bearing direction. Consequently, reproducing this wavy collagen arrangement is crucial when designing scaffolds that aim to mimic native ligament mechanics and support functional tissue regeneration.

Tissue engineering (TE) has emerged as a promising strategy to overcome the limitations of traditional grafts by combining cells, biomaterials, and bioactive cues to regenerate functional tissues. In ligament TE, a key challenge is to design scaffolds that recapitulate both the anisotropic mechanics and the microstructural organization of the native tissue. Early efforts relied on braided or electrospun scaffolds to produce aligned fibrous structures [12–18]. These systems successfully promoted cell alignment and ECM deposition. Electrospinning, in particular, enables the creation of fibrous constructs with hierarchical organization similar to native collagen bundles. However, the random nature of fiber deposition and the limited control over parameters such as spacing, crimp geometry, and pore interconnectivity remain major limitations of this method. In addition, electrospinning uses organic solvents, which raises concerns regarding cytotoxicity, solvent residues, and regulatory compliance for biomedical applications.

Structural biomimicry has therefore become a central design principle in ligament TE. By mimicking the hierarchical arrangement of collagen fibers, scaffolds can not only reproduce ligament-like mechanical responses but also provide topographical cues that direct cell alignment and differentiation [19–21]. Recent strategies increasingly focus on reproducing the hierarchical structure of the ligament to achieve functional regeneration [3]. The current challenge lies in developing fabrication approaches capable of achieving this level of architectural precision while maintaining mechanical integrity and suitable elasticity.

Additive manufacturing (AM) has opened new avenues for designing highly controlled and reproducible scaffolds for tissue regeneration. Compared to conventional fabrication techniques, AM enables precise, layer-by-layer control over scaffold geometry and internal architecture, allowing the design of structures with tailored mechanical and biological properties [22–24]. Among AM techniques, melt electrowriting (MEW) has gained particular attention for fibrous tissue applications. MEW combines principles of electrospinning and 3D printing, using an electric field to precisely guide a molten polymer jet, producing continuous micro-to submicrometer fibers with precise spatial control [25–28].

This level of precision makes MEW especially attractive for mimicking fibrous tissues such as tendons and ligaments, where the micro-architecture directly determines mechanical function. MEW can reproduce aligned, woven, or crimped fiber patterns that closely resemble native collagen arrangements [26,29]. Polycaprolactone (PCL) is commonly used in MEW because of its low melting temperature, thermal stability, and excellent processability. However, the relatively high stiffness and limited elasticity of PCL can restrict its use in applications requiring greater resilience, such as ligament or tendon regeneration. For instance, Petrigliano et al. reported that although a PCL-based scaffold implanted *in vivo* showed improved mechanical properties over time, it still did not match the mechanical performance expected for native ligament [30]. Nevertheless, the architecture of the scaffold strongly influences its final mechanical behavior and can partly compensate for material limitations [26,31,32].

Despite recent advances, few studies have systematically investigated how the micro-architecture of MEW scaffolds, particularly the waviness of fibers, affects the mechanical and biological behavior of ligament cells. The ability to control and reproduce such crimped geometries could provide critical insights into the structure–function relationships governing ligament regeneration.

The present study aims to explore the potential of biomimetic MEW scaffolds with tunable wavy architectures for ligament tissue engineering. We investigated the potential of using biomimetic design with MEW to fabricate scaffolds specifically tailored for ligament tissue regeneration. Using MEW, we fabricated fibrous scaffolds with controlled wave amplitudes and lengths to mimic the hierarchical crimp of native collagen ligament bundles. We assessed how these design variations influenced the scaffolds mechanical performance under tensile loading and proceeded by examining human ACL cell responses, including adhesion, proliferation, and expression of ligament-associated genes. Through this combined mechanical and biological analysis, our work demonstrates the potential of MEW to create structurally and mechanically biomimetic scaffolds for ligament regeneration, paving the way toward engineered constructs that more closely replicate the architecture and function of native ligaments.

## Materials and methods

### Fabrication of fibrous scaffolds

Fibrous scaffolds were fabricated using a commercially available MEW device (Spraybase, A-1204-0001-01D). Medical-grade poly(ε-caprolactone) (PCL, PC12, Mw = 120 kg.mol^−1^, Corbion) was used to produce 40-layer scaffolds. Printing parameters were set as follows: temperature 120 °C, pressure 0.15 bar, printing speed 10 mm/s, applied voltage 3.5 kV, and a nozzle-to-collector distance of 4 mm. A stainless-steel nozzle with an inner diameter of 0.25 mm was used.

The fiber spacing was fixed at 0.5 mm. Three scaffold designs were produced: a control with straight and parallel fibers (CTRL), and two wave-patterned designs with sinusoidal fibers, with amplitudes of 1 mm and periods of either 4 mm (A1P4) or 2 mm (A1P2).

The morphology of the produced scaffolds was observed and imaged through a Stereomicroscope (Nikon SMZ25). Samples were then coated with a nanometric gold layer using a Cressington 108 Auto sputter coater and imaged by Scanning Electron Microscope (SEM) using a Jeol JSM-IT200 InTouchScope in secondary electron mode, with a typical accelerating voltage of 10 kV.

### Mechanical properties

Uniaxial tensile tests were performed in displacement control using an Electroforce 3200 Series III mechanical tester (TA Instruments) equipped with a 45 N load cell. The tests were controlled and recorded via WinTest® 7 software. Samples (n = 5 per scaffold type) were stretched at a strain rate of 10% s^-1^ to the initial gauge length, up to a maximum displacement of 100 mm or until failure. This strain rate was selected to approximate the physiological loading conditions experienced by tendons during activities such as walking, as previously reported by Wren et al. [33].

To evaluate fatigue behavior, cyclic tensile tests, in displacement control, were performed at 1 Hz for 100 cycles. Strain levels for cyclic loading were selected based on the stress–strain curves, beyond the toe region: 10% strain for CTRL and A1P4 scaffolds, and 25% strain for A1P2 scaffolds. Cycles were applied at a frequency of 1 Hz for a total of 100 cycles. Each condition was tested on n = 4 specimens per scaffold type.

Mechanical data were processed to extract both apparent and net mechanical properties, following the approach described by Sensini et al. [34]. Apparent stress–strain curves were obtained by normalizing the load over the total cross-sectional area of the sample. To avoid the contribution for internal scaffold porosity, net stress was calculated by dividing the apparent stress by the scaffold volume fraction (v),computed as:

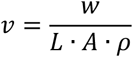

where *w* is the sample weight, *L* the length, *A* the cross-sectional area, and *ρ* the material density. A density of 1.2 g/cm^3^ was used for all calculations. From the stress–strain curves, the Young modulus (*E*), yield stress (*σ*_*Y*_), failure stress (*σ*_*F*_), permanent deformation, work to yield (*Wy*) and work to failure (*W*_*f*_) were determined. The work to yield and work to failure were obtained by integrating the area under the force–displacement curve up to the yield and failure points, respectively, using trapezoidal numerical integration. Using cyclic test data, the area under the loading and unloading curves was computed using trapezoidal integration to quantify hysteresis.

### Cell culture and seeding

Human ACL cells were obtained from patients undergoing full ligament replacement after traumatic injury, with written informed consent (METC number 2017-0183). Tissue samples were collected immediately post-surgery, placed in phosphate-buffered saline (PBS), and dissected into 2 × 2 mm pieces under an optical microscope. Surrounding adipose tissue was carefully removed and discarded. The dissected tissue fragments were washed thoroughly in PBS, transferred to 15 mL centrifuge tubes, and enzymatically digested for 4 hours at 37 °C and 5% CO2 in a solution of 0.4% collagenase (Sigma Aldrich) and 0.25% trypsin (Sigma Aldrich) prepared in PBS. Following digestion, the suspension was centrifuged at 500 rpm for 5 minutes. The supernatant containing released cells was filtered through a 0.2 µm cell strainer and diluted to a total volume of 7 mL in high-glucose Dulbecco’s Modified Eagle Medium (DMEM, 4.5 g/L glucose, Gibco) supplemented with 10% fetal bovine serum (FBS, o48B16, VWR) and 1% penicillin–streptomycin (pen/strep). Cells were seeded in 25 cm^2^ culture flasks and incubated under standard culture conditions. The culture medium was refreshed every second day until cells reached 80% confluence. Cells were passaged using 0.5% trypsin and reseeded at 10,000 cells/cm^2^. All experiments were performed using cells at passage 3 to avoid de-differentiation.

The scaffolds were clamped in CellCrown™ inserts (Z742381) to facilitate handling and ensure stability during cell culture. Sterilization was performed by immersing the scaffolds in three consecutive baths of 70% ethanol. To prevent cell attachment to the plastic surface, 24-well plates used for seeding were treated with Anti-Adherence Rinsing Solution (Stemcell Technologies) according to the manufacturer instructions. Cells were suspended in a 10% (w/v) dextran (T-500 Dextran, Pharmacosmos) solution prepared in cell culture medium following a protocol previously developed by Cámara-Torres et al. for enhanced seeding efficiency on 3D scaffolds [35]. Cells were then seeded onto the scaffolds at a density of 85,000 cells per scaffold. After 24 hours, the scaffolds were transferred to new wells, and the cell culture medium was refreshed three times per week.

### Morphological Analysis of Cell-Cultured Scaffolds

The morphology of the cell-cultured scaffolds was analyzed using SEM. Cell-containing samples were dehydrated through a graded ethanol series in water, with 15-minute incubations in 30%, 50%, 70%, 80%, 90%, 96%, and 100% ethanol solutions. Samples were then briefly air-dried and incubated sequentially in a 1:1 solution of ethanol and hexamethyldisilazane (HMDS, Sigma-Aldrich), followed by 100% HMDS, each for 15 minutes. After HMDS removal, samples were left to air-dry overnight.

For high-resolution imaging, dried samples were sputter-coated with a nanometric gold layer (Cressington 108 Auto) and imaged using the same SEM setup (JEOL JSM-IT200 InTouchScope) in secondary electron mode, with an accelerating voltage of 7 kV.

### DNA quantification

Samples were harvested on days 3, 7, 14, and 21 of culture and stored at −80 °C until analysis. Samples underwent three freeze–thaw cycles by alternating immersion in liquid nitrogen and thawing at 56 °C in a thermomixer. Each sample was then incubated overnight at 56 °C with 1 mg/mL proteinase K (Sigma Aldrich) in a buffer composed of 50 mM Tris, 1 mM EDTA, and 1 mM iodoacetamide to ensure complete cell lysis. Following digestion, samples underwent an additional three freeze–thaw cycles using the same procedure. DNA content was quantified using the CyQuant Cell Proliferation Assay Kit (ThermoFisher), according to the manufacturer’s instructions. Briefly, cellular RNA was degraded by incubating the samples at room temperature for at least 1 hour in lysis buffer containing RNase A (Thermo Fisher, diluted 1:500). A DNA standard curve was prepared as described in the kit protocol. For fluorescence measurement, 100 µL of each sample (in duplicate) was transferred to a black 96-well microplate, followed by the addition of 100 µL of 2× GR-dye solution. After mixing, the plate was incubated in the dark at room temperature for 10 minutes. Fluorescence was measured using a Clariostar plate reader (BMG Labtech) at an emission wavelength of 520 nm.

### Metabolic Activity

Cell metabolic activity was assessed using the PrestoBlue® Cell Viability Reagent (Thermo Fisher Scientific). At selected time points (days 3, 7, 10, 14, 18, and 21), culture medium was removed, and samples were gently washed twice with PBS. A working solution of PrestoBlue® (10% v/v in fresh culture medium) was freshly prepared and protected from light. A total of 100 μL of PrestoBlue® solution was added to each scaffold-containing well, along with appropriate controls: cell-free scaffold controls to account for potential background fluorescence.

Samples were incubated at 37 °C for 45 minutes in the dark. After incubation, 100 μL of the supernatant was transferred to a black 96-well plate with a transparent bottom. Fluorescence was measured using a microplate reader (excitation: 560 nm, emission: 590 nm). Background fluorescence from scaffold-only controls was subtracted from all measurements.

### Immunofluorescence

Samples were fixed by immersion in 4% paraformaldehyde (PFA) for 15 minutes, followed by rinsing with PBS and permeabilization in 0.1% Triton X-100 for 30 minutes. After fixation and permeabilization, samples were blocked overnight at 4 °C in PBS containing 1% bovine serum albumin (Roth), 0.05% TWEEN® 20 (Sigma-Aldrich), and 5% donkey serum (Abcam).

Samples were then incubated overnight at 4 °C with mouse anti-collagen I (Abcam, ab6308) and rabbit anti-tenascin-C (Bioss, BS-1039R), both diluted 1:100 in washing buffer (blocking buffer diluted 1:5 in PBS). Following primary antibody incubation, samples were washed and incubated for 1 hour with secondary antibodies: donkey anti-mouse and donkey anti-rabbit both diluted 1:200 in washing buffer. Simultaneously, Alexa Fluor™ Phalloidin 567 (Thermo Fisher Scientific) was added at a 1:100 dilution and incubated for 2 hours at room temperature.

After additional washes, samples were stained with DAPI for 30 minutes at room temperature, rinsed, and stored in the dark at 4 °C until imaging. Imaging was performed using a Leica TCS SP8 STED confocal microscope.

### Visualization and quantification of glycosaminoglycans (GAGs)

Glycosaminoglycans (GAGs) were visualized by Alcian Blue (A5268, Sigma-Aldrich) staining on scaffolds cultured for 21 days. Fixed samples (4% PFA for 15 minutes) were first immersed in 3% acetic acid for 5 minutes, then stained with Alcian Blue solution (10 g/L in 3% acetic acid, pH 2.5) for 15 minutes. After staining, samples were rinsed in running tap water for approximately 2 minutes, dipped briefly in distilled water, and counterstained with nuclear fast red for 10 minutes to label cell nuclei. Images of stained scaffolds were acquired using a stereomicroscope (Nikon SMZ25).

Quantification of GAGs was performed on the same samples used for DNA analysis, following the proteinase K digestion step. A 1,9-dimethylmethylene blue (DMMB) assay was used, prepared by dissolving 16 mg of DMMB in 5 mL of ethanol. Chondroitin sulphate from shark cartilage (Sigma-Aldrich) was used to generate the standard curve. Briefly, 150 µL of DMMB solution was mixed with 50 µL of the sample and 5 µL of 2.3 M NaCl in a black 96-well microplate. Absorbance was measured at 525 and 595 nm using a Clariostar plate reader (BMG Labtech), and the difference in absorbance was used to calculate the GAG content.

### Collagen visualization and quantification

Collagen deposition was assessed using Picrosirius Red staining (ab150681, Abcam), following the manufacturer’s instructions. Scaffolds cultured for 21 days were fixed in 4% PFA for 15 minutes, then immersed in Picrosirius Red solution for 30 minutes. Samples were subsequently rinsed twice with 0.5% acetic acid solution, followed by one rinse with absolute ethanol. Two additional dips in absolute ethanol were performed to ensure complete dehydration. Collagen fibers stained with Picrosirius Red were imaged under polarized light using an inverted Nikon Ti-S/L100 microscope equipped with a Nikon DS-Ri2 camera. Birefringent collagen structures were visualized by placing the polarized light filter at 90° relative to the analyzer. Illumination was provided by a CoolLED pE100 system for diascopic white light, and images were acquired in brightfield mode. The samples were also imaged in reflection mode at 488 nm using a Leica TCS SP8 STED confocal microscope, to visualize unstained collagen fibers based on previous works [36,37].

### PCR

To investigate the gene expression on days 7 and 21, scaffolds were washed with PBS and RNA was extracted from samples using a Maxwell RSC simplyRNA Tissue Kit with a Maxwell RSC Instrument. Concentration and purity of RNA were measured using a Spectramax QuickDrop Micro-Volume spectrophotometer. cDNA was synthetized using iScript cDNA synthesis kit (Bio-Rad) following manufacturer’s instructions. Finally, qPCR reactions were conducted in 20 μl total volume mixing cDNA and selected primers (Table 1) with SYBR Green Supermix (Qiagen). A CFX Connect Real-Time System (Bio-Rad) was used with the following thermal cycle: 95 °C for 3 min, 40 cycles for 15 s at 95 °C and 30 s at 55 °C. For each target gene, GAPDH was selected as the housekeeping gene for normalization and the 2− ΔΔCt method was performed to calculate the relative expression. Further normalization was done with reference to the relative expression of cells cultured in 2D in maintenance medium.

**Table 1.**
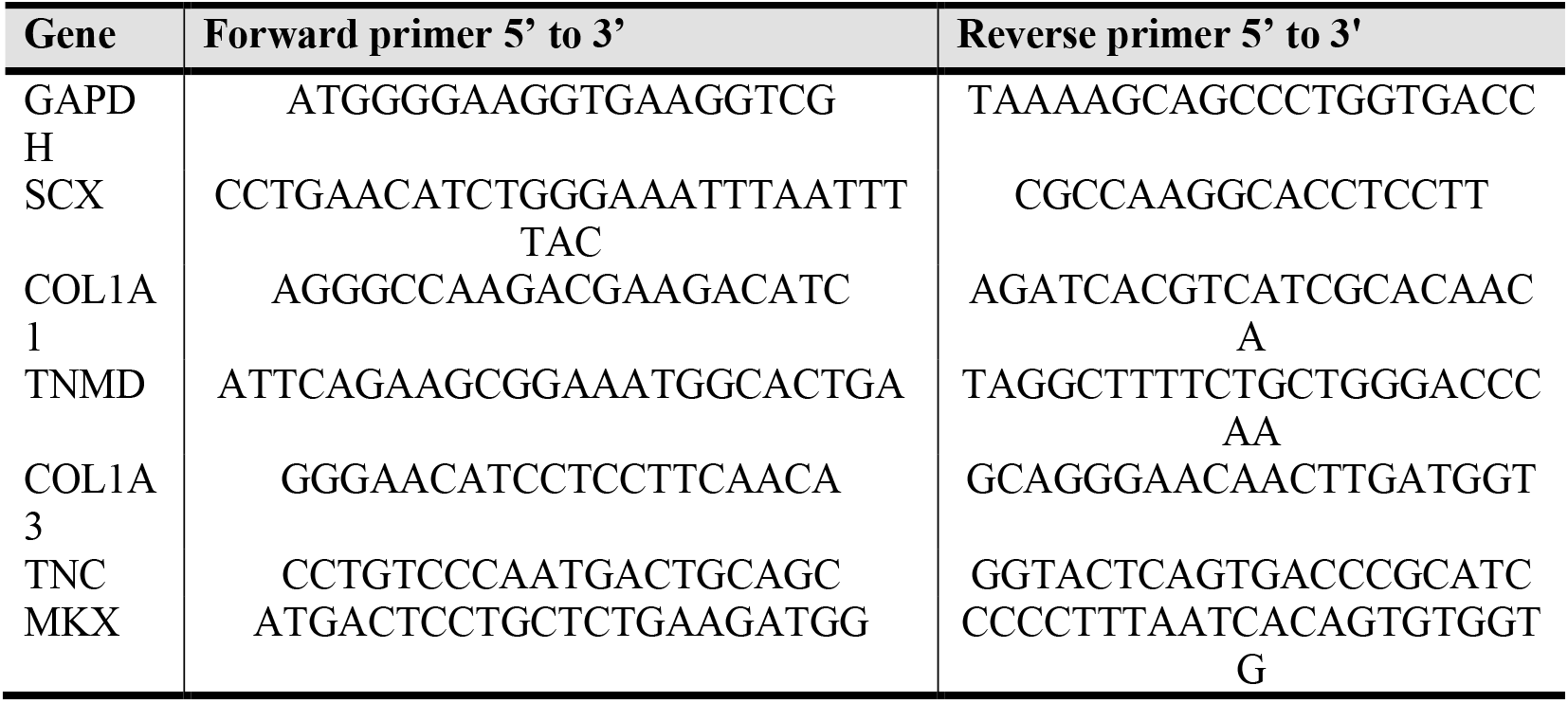
Primers used for PCR.

### Statistical analysis

Mechanical test data were analyzed using a two-way ANOVA to assess the effects of scaffold type and treatment condition (before vs. after cycling), followed by a Bonferroni post hoc test for multiple comparisons. GAG quantification and gene expression data were assessed for normality using Shapiro–Wilk and Kolmogorov–Smirnov tests, which confirmed normal distributions. Statistical comparisons between groups for GAGs were performed using a one-way ANOVA followed by a Tukey post hoc test, while gene expression data were analyzed using a two-tailed Student’s t-test. All analyses were performed using GraphPad Prism (version X, GraphPad Software).

## Results & discussion

### Scaffolds fabrication

MEW enabled the fabrication of highly regular and well-aligned PCL microfibrous meshes, demonstrating excellent control over fiber deposition and architecture (Figure 1). The printed scaffolds exhibited a consistent and reproducible pattern, with no fiber fusion or significant defects, indicating stable and optimized process parameters. The successful fabrication of wavy scaffolds further underscores the potential of this technique for producing highly organized architectures tailored to ligament tissue engineering.

**Figure 1.**
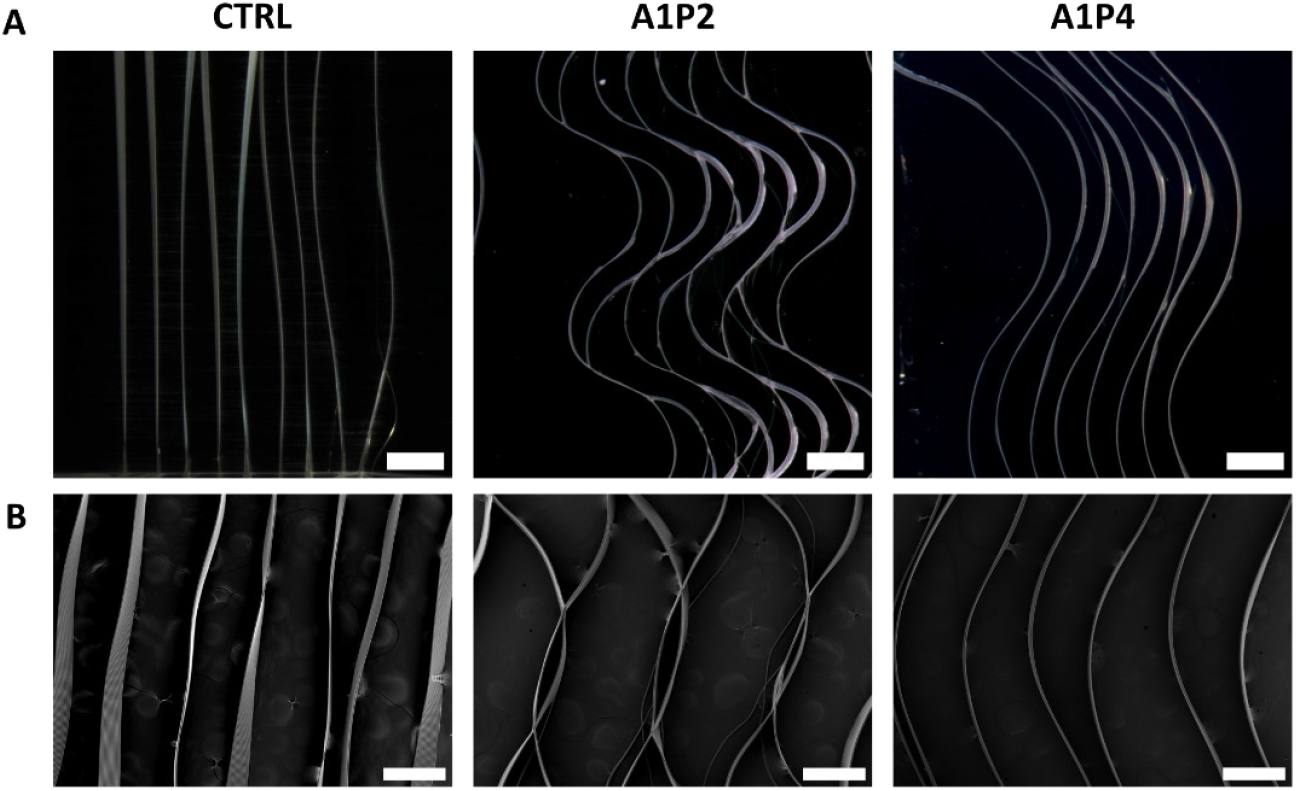
PCL scaffolds with wave-patterned architectures fabricated by MEW. Wave designs were printed with a fixed amplitude of 1 mm and varying periods of 2 mm and 4 mm. (A) Images were acquired using a stereomicroscope (Scale bar = 1 mm) and (B) a SEM (scale bar = 500 µm).

Notably, while most studies rely on the critical translation speed (CTS) to induce controlled buckling of extruded fibers and generate sinusoidal or crimped architectures [25,38], this strategy generally requires low print speeds, typically below 1 mm/s, resulting in long fabrication times. In contrast, we took advantage of the high movement resolution of our MEW system to directly print predefined wave shapes without relying on CTS-induced instabilities. This approach enabled rapid scaffold fabrication at 10 mm/s, nearly ten-fold faster than CTS-based methods, while maintaining structural fidelity. By overcoming the constraints of CTS-based printing, our method offers a more efficient and reproducible alternative for generating functionally relevant fiber geometries.

### Mechanical characterization

Mechanical characterization revealed that the scaffold architecture had a pronounced impact on tensile properties (Figure 2). CTRL scaffolds with straight fibers exhibited an apparent Young’s modulus (*E*_*A*_) of 1.25 ± 0.35 MPa and a yield stress (*σ*_*Y*_) of 0.09 ± 0.02 MPa (Figure 2B).

**Figure 2.**
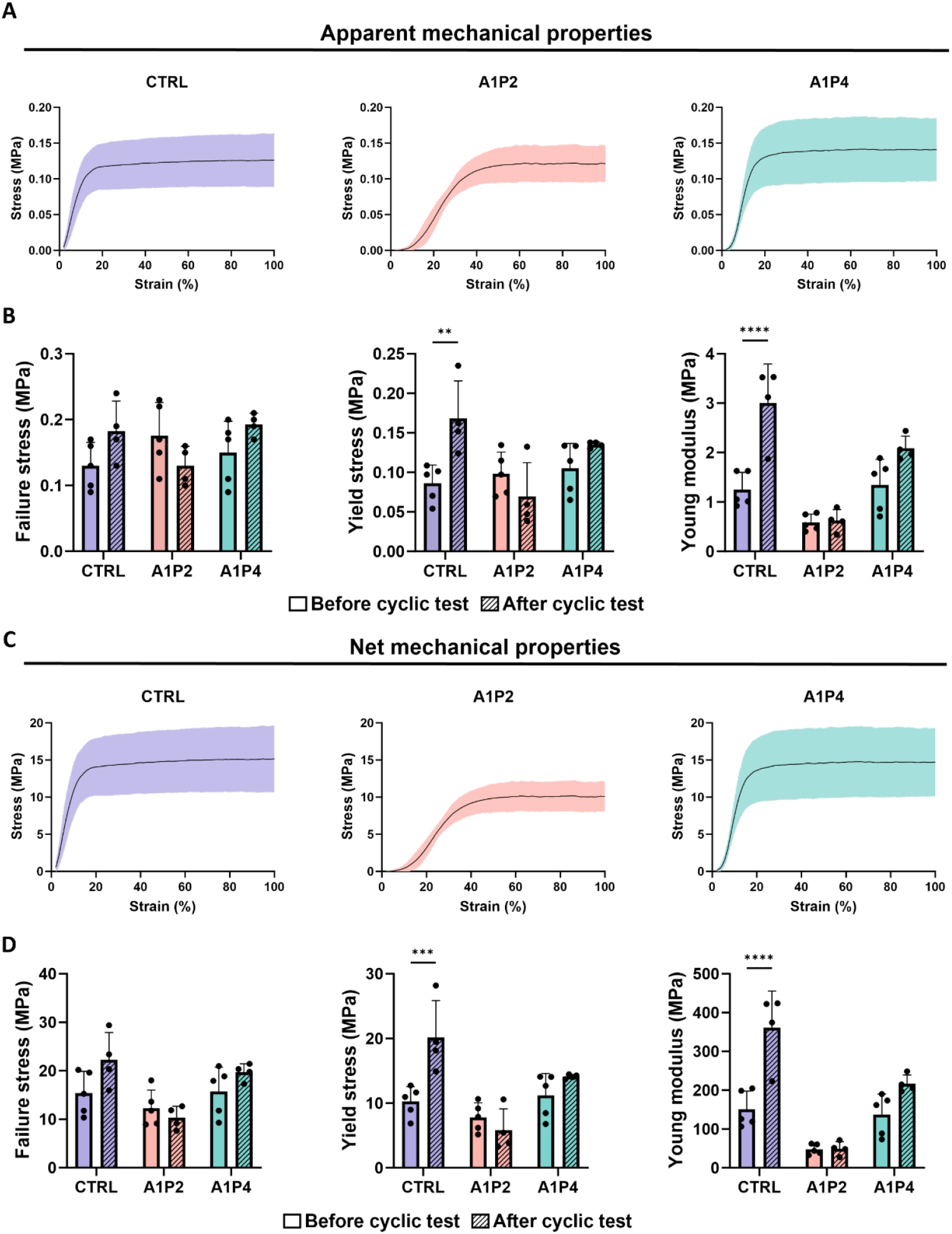
Mechanical behavior of scaffolds under uniaxial tensile loading, before and after cyclic testing. (A) Apparent stress–strain curves (mean ± SD) for each scaffold type: control with straight fibers (CTRL, mauve), short-period waves (A1P2, coral), and long-period waves (A1P4, teal). (B) Macroscopic mechanical properties: failure stress, yield stress, and Young’s modulus before (solid bars) and after (striped bars) cyclic loading. (C) Net stress–strain curves (mean ± SD) for each scaffold type. (D) Net mechanical properties recalculated based on the effective load-bearing cross-section. Data are shown as mean ± SD, individual data points represent replicates. Statistical significance: *p < 0.05, **p < 0.01, ***p < 0.001, ****p < 0.0001.

Scaffolds with short wave periods (A1P2) showed the lowest apparent modulus (*E*_*A*_ = 0.59 ± 0.17 MPa) and a yield stress of 0.10 ± 0.03 MPa, confirming their highly compliant nature and early plastic deformation (Figure 2B). In contrast, A1P4 scaffolds with longer wave periods displayed higher apparent modulus (*E*_*A*_ = 1.34 ± 0.52 MPa) and yield stress (0.11 ± 0.03 MPa), indicating greater mechanical stability (Figure 2B).

Prior to cyclic loading, A1P4 scaffolds exhibited the highest work to failure (*Wf* = 58.38 ± 20.74 mJ), consistent with their superior load-bearing capacity and mechanical robustness (Figure S2). CTRL and A1P2 scaffolds showed slightly lower but comparable *Wf* values (50.47 ± 13.80 mJ and 44.18 ± 23.94 mJ, respectively), indicating similar overall energy absorption under monotonic loading. In contrast, work to yield (*Wy*) values revealed a different trend: A1P2 exhibited the highest *Wy* (0.62 ± 0.47 mJ), suggesting that this more compliant design accommodated greater deformation before yielding (Figure S2). A1P4 and CTRL showed lower *Wy* values (0.37 ± 0.14 mJ and 0.19 ± 0.08 mJ, respectively), reflecting a lower threshold for plastic deformation.

During cyclic loading testing, the effect of the pattern/design on the mechanical properties became evident (Figure 2, Supplementary Videos S1–S3). CTRL scaffolds exhibited a marked increase in apparent modulus, reaching 3.01 ± 0.78 MPa, along with a rise in yield stress (*σ*_*Y*_ to 0.17 ± 0.05 MPa), indicative of strong viscoelastic preconditioning. A1P4 also showed a significant increase in apparent modulus (*E*_*A*_ = 2.09 ± 0.25 MPa) and yield stress (*σ*_*Y*_ = 0.13 ± 0.00 MPa), consistent with progressive fiber alignment under loading. In contrast, A1P2 remained the most compliant (*E*_*A*_ = 0.62 ± 0.22 MPa) and exhibited a slight decrease in yield stress (0.07 ± 0.04 MPa), suggesting limited structural adaptation and a reduced resistance to plastic deformation under repeated loading. Failure stress showed minimal variation post-cycling, increasing slightly for CTRL and A1P4 and decreasing for A1P2, indicating good structural resilience.

Following cyclic testing, all scaffolds exhibited a marked reduction in both *Wy* and *Wf*, reflecting fatigue-induced degradation of their load-bearing capacity (Figure S2). The drop in work to failure was significant across all groups, decreasing by about 80 % relative to pre-cyclic values (CTRL: 10.80 ± 16.02 mJ; A1P2: 7.82 ± 4.00 mJ; A1P4: 6.26 ± 7.51 mJ). This substantial loss is consistent with the progressive softening and microstructural rearrangements observed under repeated loading. In contrast, changes in work to yield were more moderate. CTRL showed a slight increase (from 0.19 ± 0.08 mJ to 0.27 ± 0.09 mJ), suggesting limited adaptation or preconditioning, while A1P2 and A1P4 decreased to 0.39 ± 0.67 mJ and 0.18 ± 0.06 mJ, respectively, indicating reduced elastic–plastic energy absorption. Among the tested designs, CTRL retained the highest post-cyclic *Wy* and *Wf* values overall, but with large variability, whereas A1P4 showed more consistent behavior, suggesting better fatigue tolerance despite its higher apparent modulus.

To better reflect the intrinsic material contribution, mechanical properties were normalized to the effective load-bearing cross-section (Figure 2C-D). Net values for Young Modulus, yield, and failure stress were nearly one order of magnitude higher than macro-scale values. This difference highlights how scaffold porosity lowers the overall measured apparent modulus and strength, even though the fiber material itself is much stronger. This phenomenon has already been documented for porous and additively manufactured materials [34,39–41]. Net values are also more biologically relevant, as cells adhere directly to individual fibers and therefore experience the local (net) stresses rather than the apparent stresses calculated over the porous scaffold. Apparent properties remain useful for comparison with native ligament, which is not porous, but they can underestimate the stiffness actually perceived by cells. Importantly, the relative differences among scaffold architectures were preserved: scaffolds with short wave patterns (A1P2) remained the most compliant, while those with longer wave periods (A1P4) exhibited the highest net modulus and strength. This confirms that the observed mechanical differences primarily arise from geometry rather than material composition.

Cyclic loading analysis further highlighted architecture-dependent responses (Figure 3A, Figure S2). All scaffolds exhibited a sharp reduction in hysteresis between the first and second cycles (Figure 3B, Figure S2), followed by gradual stabilization by Cycle 100. CTRL scaffolds consistently displayed the highest hysteresis values (0.09 ± 0.01 at Cycle 1 to 0.04 ± 0.00 at Cycle 100), indicating greater energy dissipation but also steady stabilization. A1P4 started lower (0.06 ± 0.04 at Cycle 1 to 0.03 ± 0.02 at Cycle 100) and stabilized quickly, while A1P2 began at a similar level (0.08 ± 0.07) but with markedly higher variability across replicates, settling to 0.03 ± 0.03 by Cycle 100. This variability suggests localized differences in fiber deformation in A1P2, likely influenced by the fiber cross-sections that act as anchoring points and create localized stress concentrations (Supplementary Video S2). Permanent deformation followed a similar architecture-dependent trend (Figure 3C). A1P2 scaffolds displayed substantial and irreversible elongation, with permanent deformation reaching 19.14 ± 2.80 % by Cycle 100 (Figure 3C). This reflects a high degree of structural rearrangement under load, likely due to fiber uncrimping and plastic deformation. In contrast, CTRL and A1P4 scaffolds maintained low permanent deformation (<5%) across all cycles, indicating excellent dimensional stability under repeated mechanical stress (Figure 3C). Interestingly, despite the modest permanent deformation observed in CTRL scaffolds, macroscopic examination after testing revealed noticeable elongation, likely due to local fiber realignment not fully captured by axial strain measurements (Supplementary Video S1).

**Figure 3.**
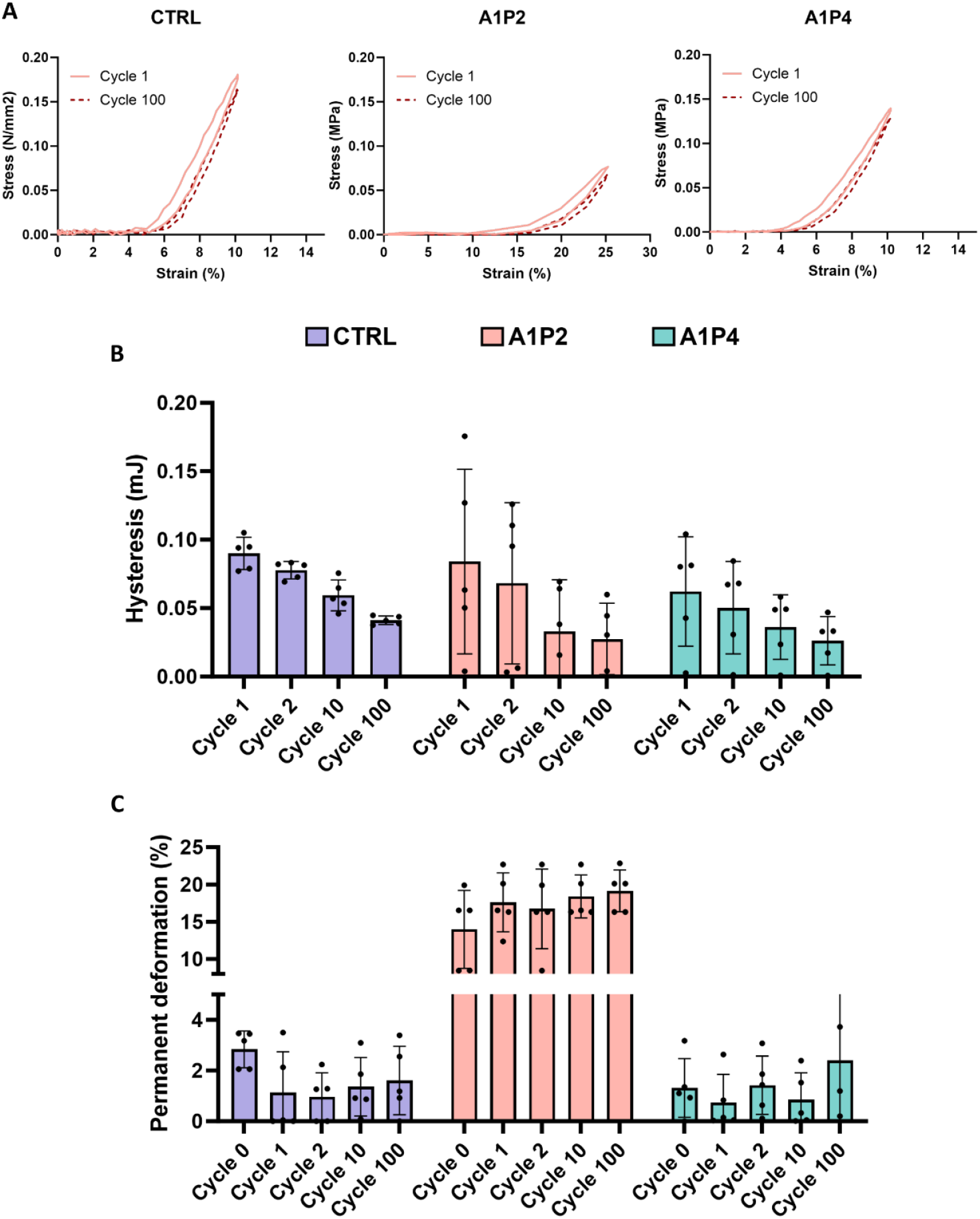
Cyclic mechanical behavior of scaffolds under repeated loading. (A) Representative stress– strain curves from Cycle 1 and Cycle 100 for scaffolds with straight fibers (CTRL), short-period waves (A1P2), and long-period waves (A1P4), illustrating structural adaptation over time. Strain levels were chosen based on the end of the toe region for each scaffold type. (B) Hysteresis energy measured over cycles. (C) Permanent deformation at selected cycles. Data are presented as mean ± SD, individual data points represent replicates.

Overall, these results highlight the strong influence of scaffold geometry on mechanical behavior under both monotonic and fatigue-like conditions. A1P4 scaffolds consistently demonstrated high elastic modulus and yield stress, while maintaining work to failure values with relatively low dispersion after cycling, indicating good fatigue tolerance. In contrast, A1P2 scaffolds maintained sufficient work to yield and work to failure to withstand early mechanical challenges while remaining significantly more compliant and prone to permanent deformation under repeated loading. Their mechanically adaptive behavior, coupled with high compliance, may support favorable cellular responses and scaffold remodeling.

These findings align with previous studies showing that scaffold architecture, particularly fiber orientation, spacing, and crimping, affects mechanical performance, independent of the base material [38,42]. Our results reinforce this structure-function relationship, showing that subtle adjustments in fiber geometry can generate distinct mechanical profiles.

Together, these results demonstrate that MEW allows precise, architecture-driven modulation of mechanical properties, including fatigue response and structural stability. Scaffolds like A1P4 offer a promising balance of elastic modulus, resilience, and repeatable behavior, while more compliant designs like A1P2 may be particularly relevant for mimicking the soft and anisotropic mechanics of the ligament.

### Cell adhesion, proliferation, and activity

For this preliminary *in vitro* investigation, we selected the A1P2 scaffold design to demonstrate the compatibility of this innovative architecture with human anterior cruciate ligament (ACL) cells. Given that the primary objective of this study was to assess cell attachment and ECM production on PCL waves, the highly compliant nature of A1P2 provided a suitable model to explore cellular behavior in a distinctive microenvironment.

From day 7 (D7), cells were clearly observed throughout the scaffold surface, aligning along the wavy walls and preferentially attaching and spreading at cross-section junctions (indicated by orange arrows in Figure 4A), likely due to increased surface area and curvature. This behavior is consistent with prior reports showing that micro-scale topographies and fiber curvature can guide cytoskeletal organization and promote cell anchorage [43]. By day 21 (D21), SEM pictures revealed dense cell coverage and partial pore filling, suggesting active colonization and progressive matrix formation across the scaffold.

**Figure 4.**
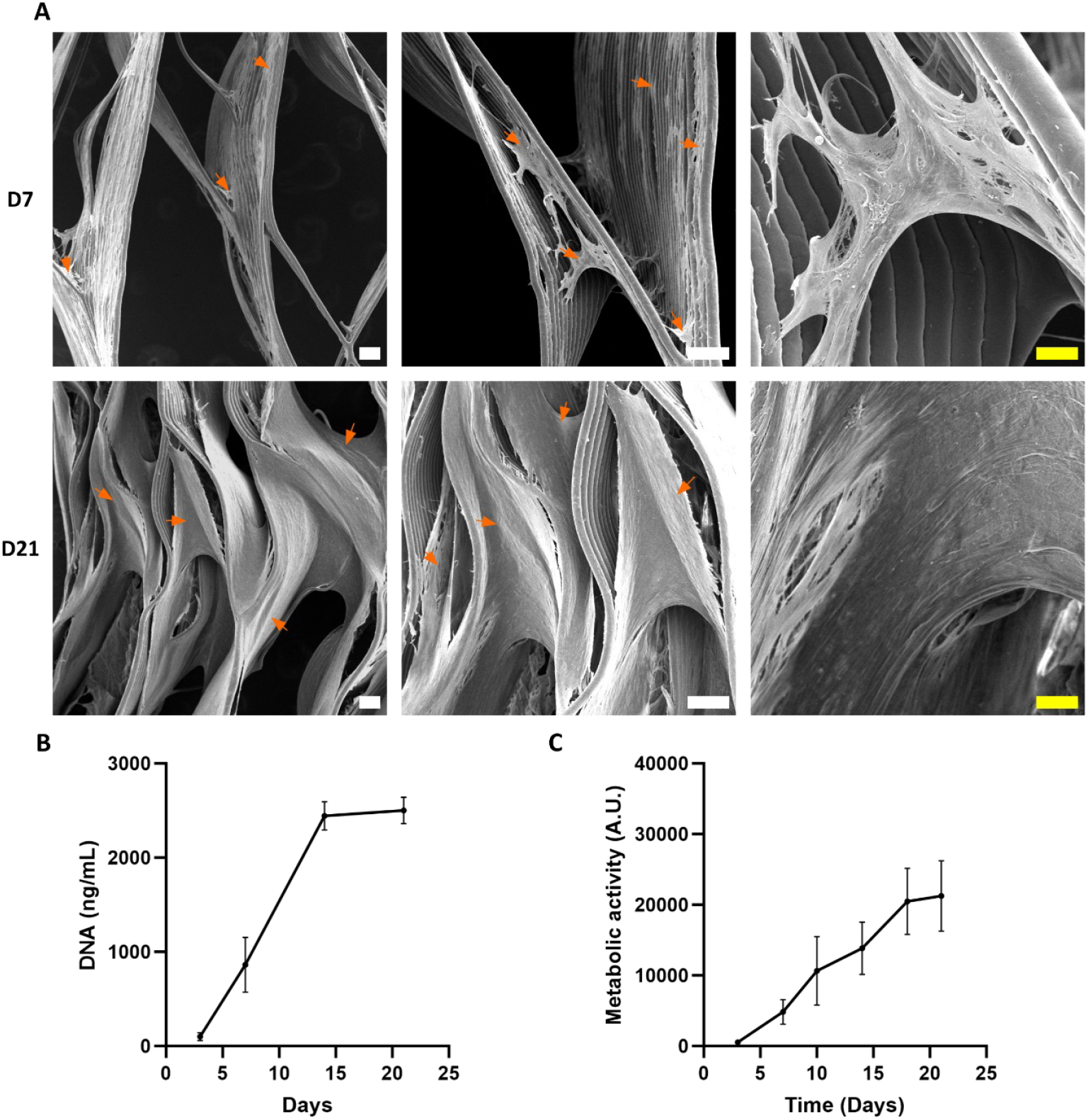
Evaluation of ACL cell colonization and activity on wavy scaffolds. (A) SEM images acquired at day 7 and day 21 to visualize cell distribution and scaffold surface coverage. White scale bars = 100 µm; yellow scale bar = 20 µm. (B) DNA content was quantified at days 3, 7, 15, and 21 to assess cell proliferation. Data represent mean ± SD from 2 independent experiments with 3 technical replicates each (n = 6). (C) Metabolic activity was measured at days 3, 7, 10, 15, 18, and 21 to evaluate cell activity and function. Data represent mean ± SD from 2 independent experiments with 5 technical replicates each (n = 10).

Quantitative DNA measurements confirmed progressive cell proliferation on the scaffolds throughout the culture period (Figure 4B). A steep increase in DNA content was observed between day 3 (D3) and day 15 (D15), indicating active cell division. From D15 to D21, DNA levels appeared to plateau, suggesting that cells had reached confluence or a state of contact inhibition. In parallel, metabolic activity, assessed throughout the experiment, increased continuously from D3 to D21, reflecting sustained cell viability and functional activity even after proliferation slowed down (Figure 4C).

Taken together, these observations support that the developed scaffolds provide a supportive microenvironment for ligament cells, enabling attachment, spreading, proliferation and matrix production.

### ECM deposition and ligament-related marker expression

To evaluate ECM production and ligament-associated marker expression, human ACL cells were cultured on scaffolds for up to 21 days and analyzed by immunostaining, histology, and biochemical assays (Figure 5).

**Figure 5.**
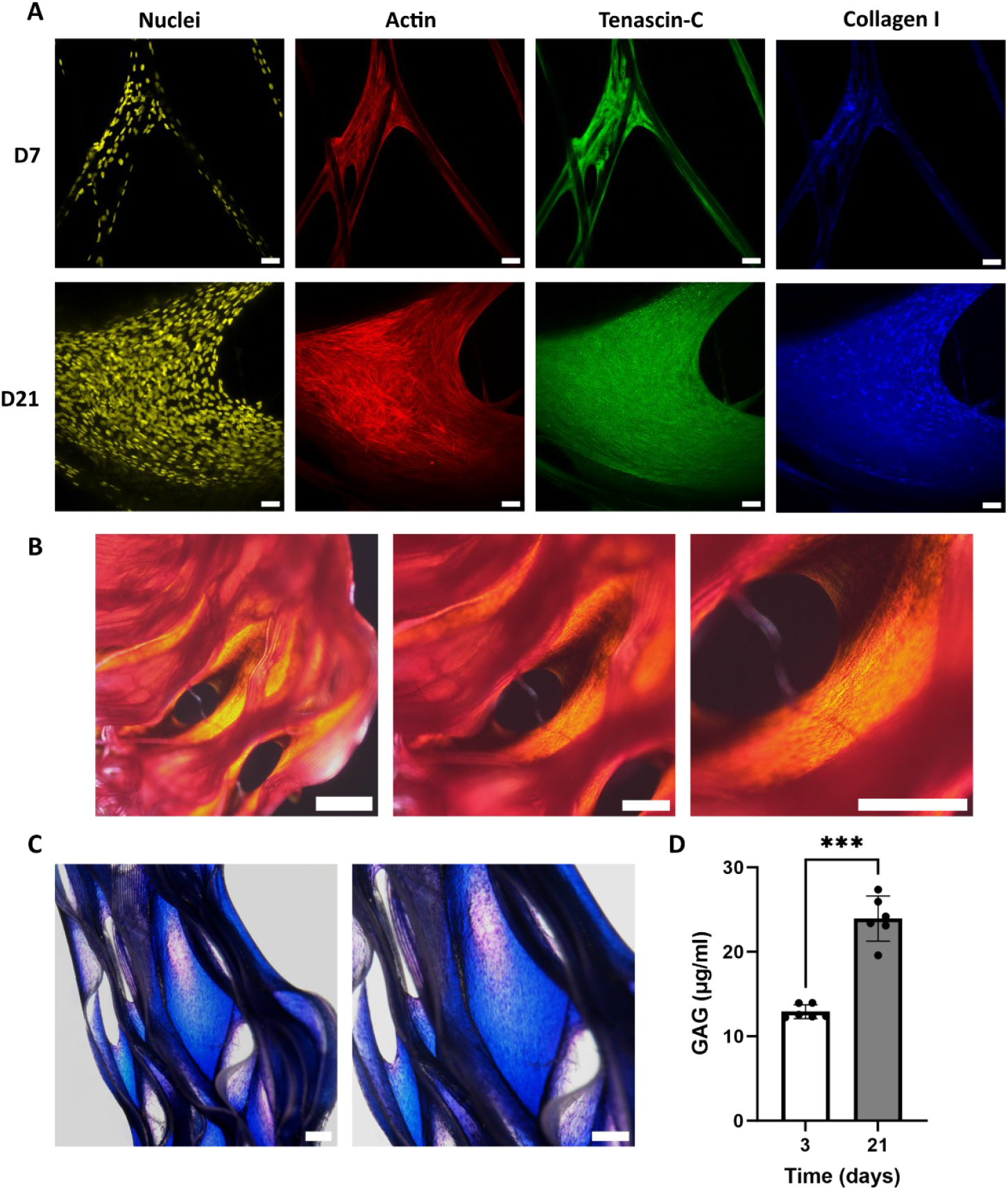
Cell colonization and matrix deposition on scaffolds seeded with human ACL cells. (A) Representative confocal images showing immunofluorescent staining at day 7 (top row) and day 21 (bottom row). Nuclei (DAPI, yellow), actin cytoskeleton (phalloidin, red), Tenascin-C (green), and collagen type I (blue) are displayed separately. Scale bar = 50 µm. (B) Representative images of Picrosirius Red staining on full construct. Scale bars = 200 µm. (C) Representative histological images of Alcian Blue staining (GAGs, blue) counterstained with Nuclear Fast Red (nuclei, pink), illustrating GAG matrix deposition at day 21. Scale bars = 200 µm. (D) Quantification of GAG production by ACL cells on A1P2 scaffolds. Data represent mean ± SD from 2 independent experiments with 3 technical replicates each (n = 6). ***p < 0.001.

At D7, actin and nuclei staining revealed early scaffold colonization, with cells aligning along scaffold walls and accumulating at crossing points (Figure 5A), which was in line with the SEM pictures at D7 (Figure 4A). By D21, cell density had increased markedly, with pronounced actin fiber organization and cytoskeletal maturation, mimicking ligament-like morphology and highlighting the influence of fiber geometry in guiding cell organization (Figure 5A). Similar alignment behavior has been reported on anisotropic scaffolds designed to reproduce ligament architecture, where cells align along patterned features in response to topographical cues [19,44]. This organization is coherent with the parallel arrangement of cells and ECM fibrils observed in native ligament-proper tissue [8,10].

Tenascin-C was detected from D7 and strongly upregulated by D21, indicating activation of ligament-associated remodeling pathways (Figure 5A). This matricellular protein is typically expressed in mechanically active connective tissues and is associated with dynamic matrix remodeling and mechanotransduction processes [45–47]. Its high expression therefore represents a positive indicator of ligament-like remodeling activity within the scaffold.

Collagen type I was already present at D7 and became more abundant and uniformly distributed by D21, indicating sustained structural ECM synthesis (Figure 5A). Collagen fiber alignment could be observed along scaffold curvature (Figure 5B, Figure S4-S5) but appeared more disorganized at fiber junctions (Figure S5). This ECM organization is consistent with the observed cell orientation, as reported in other ligament-mimetic scaffolds, where aligned architectures promote collagen alignment and matrix anisotropy [16,19,41].

The organization of ECM is crucial in load-bearing tissues like ligaments, as it directly influences their mechanical performance and biological function. Accordingly, scaffold architectural cues that guide ECM deposition and fiber alignment play a central role in recreating the anisotropic properties of native ligament tissue [48]. In this context, the wavy architecture of scaffolds demonstrates that precise microstructural control can guide both cellular and matrix organization, supporting the development of morphologically relevant ligament-like constructs.

We further proceeded by analyzing the GAGs expression on the cultured scaffolds. GAGs are of vital importance in the ACL, conferring strength and elasticity. Loss or absence of GAGs is known to reduce tendon mechanical properties [48]. Alcian blue staining confirmed the deposition of glycosaminoglycans (GAGs) within the scaffold architecture (Figure 5C). Quantification showed a significant increase in GAG content between D3 and D21 (p < 0.001) (Figure 5D), with an approximately 1.85-fold rise, reflecting progressive ECM maturation.

Together, these results demonstrate that the scaffold supported ligament cell adhesion, spreading, and progressive ECM deposition, while promoting the concurrent expression of structural (COL1) and remodeling (TNC, GAGs) markers. This combination highlights the potential of the architecture to guide early tissue formation and ligament-like remodeling.

### Gene expression

To assess the evolution of ligament-associated gene expression in ACL cells cultured on the scaffold, RT-qPCR analysis was performed at D7 and D21 (Figure 6). The temporal expression patterns reflected changes in cell activity and phenotype maintenance under static 3D culture conditions.

**Figure 6.**
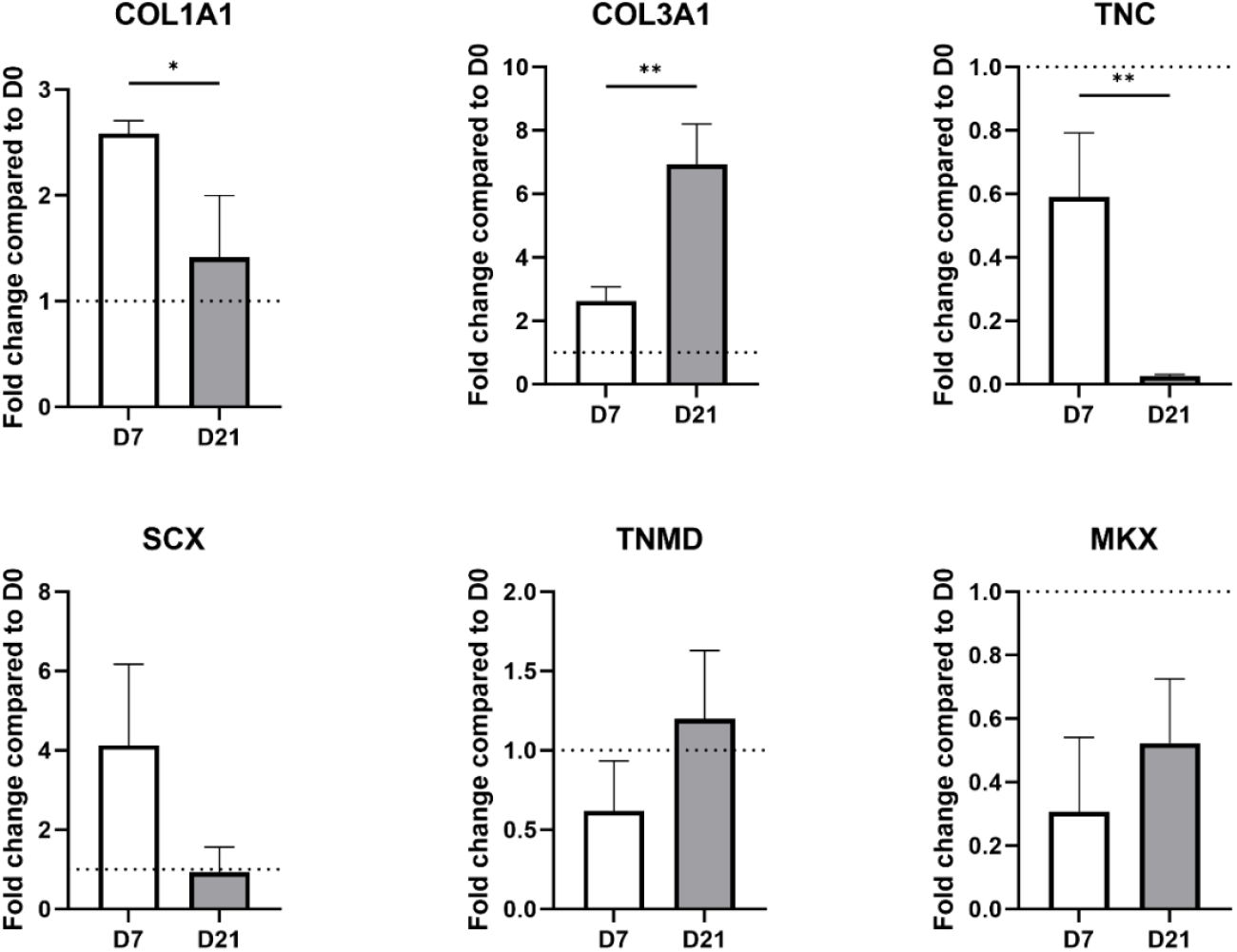
Temporal gene expression of ligament-associated markers in ACL-derived cells cultured on scaffolds. RT-qPCR analysis was performed at D7 (white) and D21 (grey) to assess the expression of key ligament-related genes. Gene expression levels were normalized to housekeeping genes and expressed relative to ACL cells cultured in 2D at day 0 (D0), set as 1 (dotted line indicates the reference). Data represent mean ± SD (n = 3); p < 0.05, p < 0.01.

COL1A1, the principal structural component of ligament ECM, was highly expressed at D7, indicating active early matrix synthesis consistent with the observed collagen deposition by immunofluorescence. Its subsequent significant downregulation at day 21 may reflect a transition from matrix production to remodeling, or a partial reduction in anabolic activity under prolonged static conditions. This decrease is consistent with previous reports showing that mechanical stimulation sustains or enhances COL1A1 expression, suggesting that its downregulation here may be linked to the absence of dynamic loading [49–52]. In an *in vitro* model, Park et al. demonstrated that scaffold-induced compression significantly increased COL1A1 expression and matrix organization in ACL-derived cells, promoting better integration and remodeling [50]. Similarly, Steffen et al. reported in an *in vivo* model that mechanical loading following ACL injury upregulated COL1A1, whereas unloading led to impaired matrix restoration [51].

COL3A1, a marker of early tissue remodeling and immature ECM, showed a significant upregulation from D7 to D21, suggesting that remodeling processes remained active. This is consistent with observations from Steffen et al., where COL3A1 remained elevated in unloaded or injured ligament, reflecting an immature or less organized matrix phenotype [51].

*Tenascin-C* (TNC) was detected at D7, but significantly reduced at D21, consistent with its transient expression during early remodeling and its mechanosensitive regulation. Reduced TNC expression likely reflects limited mechanotransductive signaling under static culture, as previously shown by Järvinen et al., where immobilization led to marked TNC downregulation in connective tissues [45].

*Scleraxis* (SCX), a key transcription factor governing tendon and ligament differentiation, was also upregulated at D7 but declined at D21. This decrease may reflect a temporal regulation of SCX expression, with an early activation phase followed by downregulation over time, as reported during tendon development and differentiation processes [53,54]. It could also indicate reduced tenogenic stimulation in the absence of mechanical cues, consistent with findings from Steffen et al., who reported better maintenance of SCX expression under mechanical loading *in vivo* [51]. Similarly, Nichols et al. showed that SCX expression is not maintained under static conditions, even when initially overexpressed, and that it remained elevated only when combined with cyclic mechanical strain in 3D culture [55]. Notably, SCX also directly regulates COL1A1, and its reduction may contribute to the observed decline in collagen expression. Nichols et al. further demonstrated that cyclic strain preserved COL1A1 expression only when SCX was overexpressed, reinforcing the link between mechanical input, transcriptional regulation, and matrix synthesis. Interestingly, *Tenomodulin* (TNMD) expression was maintained in SCX-overexpressing constructs even under strain, suggesting that some downstream ligament markers may persist even after SCX levels decline.

Among late-stage markers, *Tenomodulin* (TNMD) showed slight upregulation between D7 and D21, possibly indicating initial steps of matrix organization, while *Mohawk* (MKX) remained low at both time points. These findings suggest that while the scaffold environment supports initial ECM production, it is insufficient to mature the full ligament phenotype of ACL cells, likely due to the absence of mechanical reinforcement [56,57].

Collectively, these results demonstrate that ACL-derived cells exhibit early ligament-like activity when cultured on the scaffolds, but begin to lose phenotypic markers over time in static conditions. This underscores the importance of integrating mechanical stimulation in future studies to preserve and enhance ACL-specific features. Nevertheless, even in the absence of mechanical stimulation, the scaffolds supported robust cell organization and early matrix deposition, highlighting their strong potential for ligament tissue engineering.

This study demonstrates that architecturally tuned MEW scaffolds can effectively modulate mechanical performance and support early ligament-like cell behavior and matrix production. However, some limitations should be acknowledged. All biological experiments were conducted under static culture conditions, which likely contributed to the decline in ligament-associated gene expression over time. As ligament cells are highly responsive to mechanical stimulation, incorporating dynamic tensile loading using a bioreactor system may be essential to preserve phenotype, stimulate ECM maturation, and enhance functional tissue development. Nevertheless, investigating cell–material interactions and biological responses under static conditions represents an important step toward validating the behavior of this newly designed scaffold architecture. Understanding how cells sense and adapt to the scaffold mechanical and structural cues provides a necessary foundation for future studies integrating dynamic loading and more complex culture environments.

Moreover, further optimization may be needed to fully match target tissue properties, especially under dynamic or multidirectional loading. Nevertheless, the combination of precise MEW fabrication, geometry-driven mechanical tunability, and supportive cell responses provides a strong foundation for advancing functional ligament scaffold design.

Finally, dynamic culture is likely to enhance construct performance. Prior work has shown that bioreactor-mediated mechanical stimulation can markedly increase the tensile modulus of aligned PCL fiber scaffolds over 21 days (Banik & Brown 2020). In our context, applying controlled cyclic tension to MEW wave architectures may similarly promote matrix organization, elevate mechanical properties, and promote ligament-associated gene expression, and will be the focus of future work.

## Conclusion

This study demonstrates the feasibility of using MEW to fabricate wave-architecture PCL scaffolds with mechanical and biological properties relevant to ligament tissue engineering. By precisely tuning fiber geometry, we generated scaffold architectures with distinct mechanical behaviors. This control over architecture allowed modulation of mechanical performance and fatigue response, ranging from highly compliant to structurally resilient designs. Wave-based scaffolds supported robust ACL cell adhesion, proliferation, and early matrix production. The scaffold promoted spatially organized cell alignment and expression of key ligament-associated markers, validating its potential as a biomimetic template for early ligament regeneration. While static conditions limited long-term maintenance of the phenotype, these findings establish a solid foundation for future studies integrating dynamic mechanical stimulation to enhance ECM maturation and functional tissue development. Overall, this work underscores the potential of geometry-driven design in soft-tissue engineering and positions MEW as a versatile platform for the fabrication of next-generation ligament scaffolds with tunable mechanical and biological performance.

## Acknowledgements

This work was supported by the European Union’s Horizon 2020 research and innovation programme (grant agreement No. 953169, Interlynk). Funding for Alberto Sensini was provided by the Horizon Europe Marie Skłodowska-Curie Postdoctoral Fellowship project 3NTHESES (No. 101061826). The authors gratefully acknowledge Corbion for providing the PCL used in this project. The authors also thank Dr. Francesca Giacomini for performing the second harmonic generation imaging acquisitions.

## Notes

### Competing Interest Statement

The authors have declared no competing interest.

