## Supplementary material for "Wave-inspired MEW scaffolds for enhanced ligament tissue regeneration": Figure S1

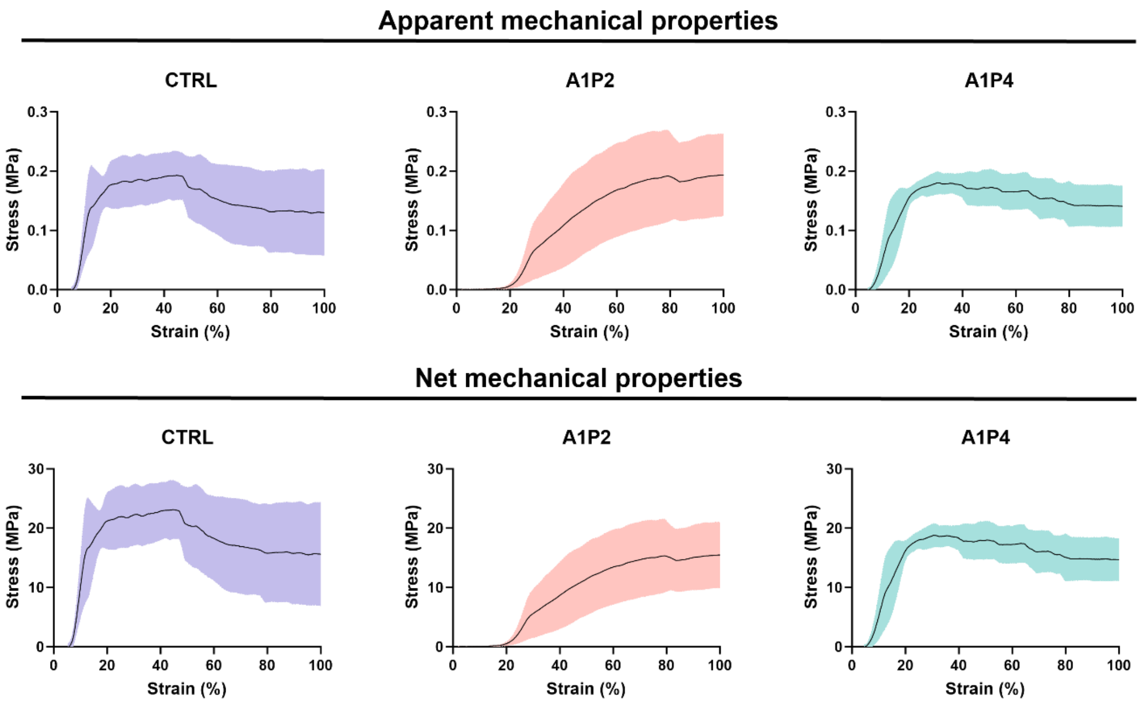


*Figure S1. Apparent and net stress–strain response of scaffolds during uniaxial extension after cyclic loading. Stress–strain curves (mean ± SD) recorded during maximal extension tests performed immediately after cyclic loading for scaffolds with straight fibers (CTRL), short-period waves (A1P2), and long-period waves (A1P4).*
