## Supplementary material for "Wave-inspired MEW scaffolds for enhanced ligament tissue regeneration": Figure S2

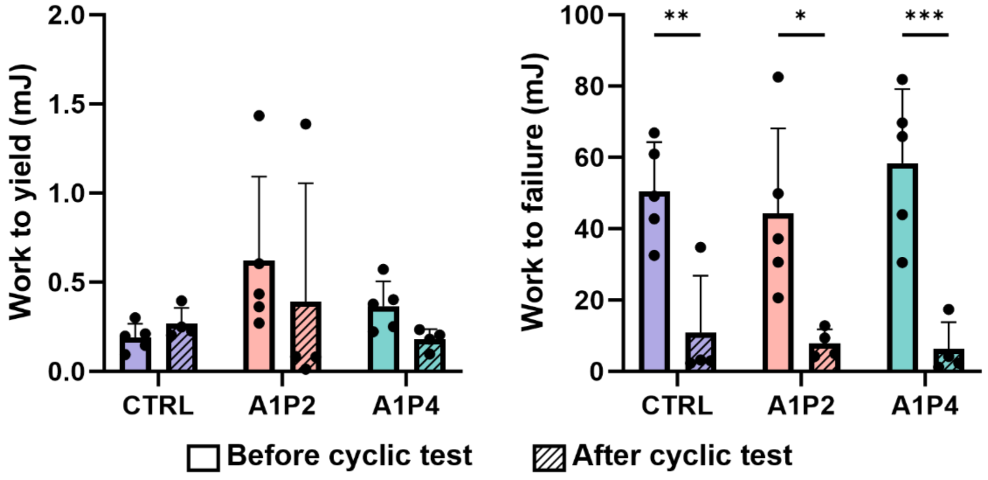


*Figure S2. Apparent work to yield and failure before (solid bars) and after cyclic testing (striped bars) for each scaffold type: control with straight fibers (CTRL), short-period waves (A1P2), and long-period waves (A1P4). Data are shown as mean ± SD, individual data points represent replicates. Statistical significance: *p < 0.05, **p < 0.01, ***p < 0.001, ****p < 0.0001.*
