## Supplementary material for "Wave-inspired MEW scaffolds for enhanced ligament tissue regeneration": Figure S3

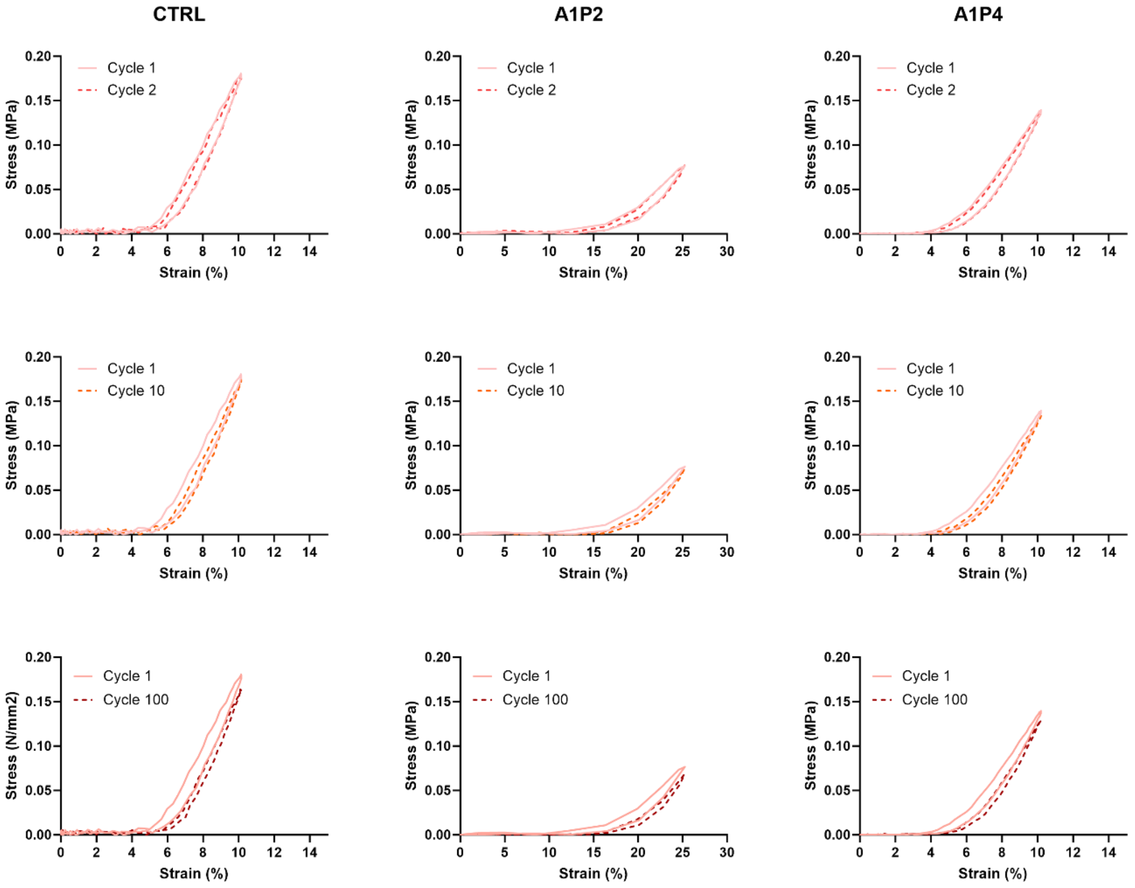


*Figure S3. Evolution of the apparent stress–strain behavior over repeated loading cycles for each scaffold architecture. Representative stress–strain curves from Cycle 1 overlaid with Cycles 2, 10, and 100 for scaffolds with straight fibers (CTRL), short-period waves (A1P2), and long-period waves (A1P4).*
