## Supplementary material for "Wave-inspired MEW scaffolds for enhanced ligament tissue regeneration": Figure S4

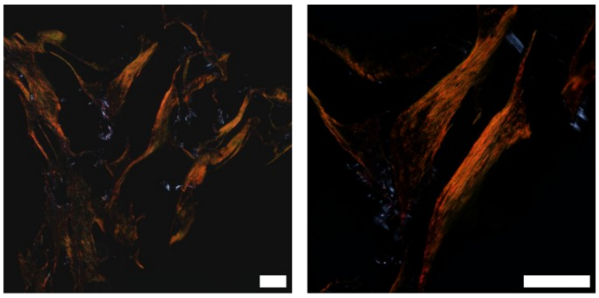

*Figure S4. Picrosirius Red staining of ACL cell–seeded scaffolds. Representative histological images illustrating collagen fiber organization at day 21. Scale bars = 200 µm.*
