## Supplementary material for "Wave-inspired MEW scaffolds for enhanced ligament tissue regeneration": Figure S5

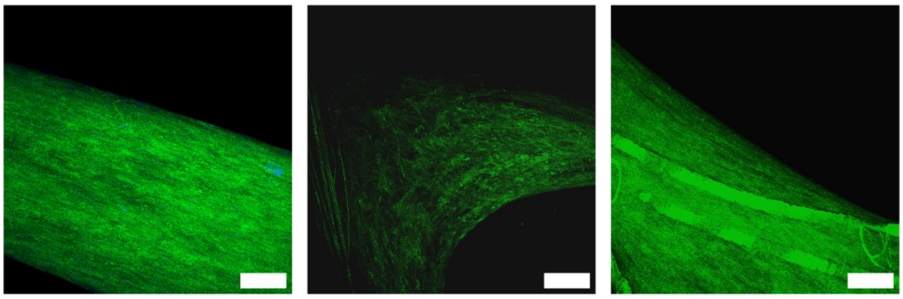

*Figure S5. Reflection-mode confocal imaging of unstained collagen fibers in ACL cell–seeded scaffolds at day 21. Samples were imaged at 488 nm using a Leica TCS SP8 STED confocal microscope. Scale bars = 200 µm.*
